# Characterization of exome variants and their metabolic impact in 6,716 American Indians from Southwest US

**DOI:** 10.1101/2020.02.21.938936

**Authors:** Hye In Kim, Nehal Gosalia, Bin Ye, Regeneron Genetics Center, Çiğdem Köroğlu, Robert L. Hanson, Wen-Chi Hsueh, William C. Knowler, Leslie J. Baier, Clifton Bogardus, Alan R. Shuldiner, Cristopher V. Van Hout

## Abstract

Applying whole exome sequencing (WES) to populations with unique genetic architecture has the potential to reveal novel genes and variants associated with traits and diseases. We sequenced and analyzed the exomes of 6,716 individuals from an American Indian population in Southwest US (Southwestern American Indian, or SWAI) with well-characterized metabolic traits. We found that individuals of SWAI have distinct allelic architecture compared to individuals with European and East Asian ancestry, with many predicted loss-of-function (pLOF) and nonsynonymous variants that were highly enriched or private in SWAI. We evaluated gene-level associations with metabolic traits using pLOF and nonsynonymous variants in SWAI. Many of the candidate genes from previous GWAS studies for body mass index, type 2 diabetes, and plasma lipid levels were associated with respective traits in SWAI. Notably, these associations were mainly driven by pLOF and nonsynonymous variants that are unique or highly enriched in American Indians, many of which have not been observed in other populations or functionally characterized. Our study illustrates the utility and potential of WES in American Indians to prioritize candidate effector genes within GWAS loci and to find novel variants in known diseases genes with potential clinical impact.

## Introduction

The genetic architecture of a population is influenced by the specific demographic history that the population has undergone. Founder and bottleneck events and subsequent reproductive isolation can result in a dramatic change in the allele frequency spectrum, potentially increasing the frequency of rare functional variants due to random genetic drift, thus allowing greater statistical power to detect the association of such variants with traits of interest^1-7^. American Indians are predicted to have gone through a series of founder and bottleneck events. One such bottleneck occurred around 15,000 years ago, when a small number of Eurasians are thought to have migrated across the Bering Strait and settled into the American continent^8^. In addition, the European colonization of the Americas led to other bottleneck events around 500 years ago^9^. Consistent with this history, American Indians have distinct genetic background compared to several cosmopolitan populations^10; 11^.

The specific population of the study consists of American Indians from the Southwestern region of the United States (SWAI). This population has very high prevalence of obesity and type 2 diabetes and has been deeply characterized for metabolic traits^12-14^. Previously, genetic studies have been conducted in this population with specific focus on metabolic traits, including genome-wide linkage analyses^15^, genome-wide association studies (GWAS)^16-20^, assessment of genes and/or variants found in GWAS studies in other ancestry groups^21-26^, and targeted sequencing of physiologic candidate genes^27-32^. These approaches have found common and rare variants that are associated with metabolic traits and disease status in this population; however, a systematic examination of coding variation across the genome and its potential impact has not been fully explored.

In this study, we sequenced the whole exomes of 6,716 American Indians and found a total of ∼1.2 million variants including 16,880 predicted loss-of-function variants and 258,306 nonsynonymous variants, many of which have not been described before. The goal of our study was to characterize the exome architecture of American Indians in comparison to cosmopolitan populations and examine the phenotypic impact of rare coding variants that are either private or enriched in this population.

## Subjects and Methods

### Study Subjects

The study participants were individuals with American Indian ancestry from the Southwestern region of the United States (referred to as “SWAI”) who enrolled in a longitudinal study described previously^14; 33^. Measurements included height and weight for body mass index (BMI) calculation and fasting lipid levels. Type 2 diabetes (T2D) status was determined based on the criteria of the American Diabetes Association or the review of the medical records. The self-reported number of great grandparents that were American Indian was recorded as a measure of admixture. Individuals with all eight American Indian great grandparents are herein referred to as “full American Indians”. DNA from blood of the participants was collected to evaluate the genetic etiology of metabolic disorders. The study protocol was approved by the Institutional Review Board (IRB) of the National Institute of Diabetes and Digestive and Kidney Diseases. Informed consent was obtained from all participants.

Individuals from two additional studies were included as references for comparison. The DiscovEHR study is a collaborative project between the Regeneron Genetics Center and Geisinger Health System based in Pennsylvania with participants who enrolled in Geisinger’s MyCode Community Health Initiative^34^. The study was approved by the IRB at the Geisinger Health System. The TAICHI study is a collaborative study with participants recruited at several academic centers in Taiwan^35^. The study was approved by the IRBs at all participating centers (Taichung Veteran’s General Hospital, Tri-Service General Hospital, the National Taiwan University Hospital, and the National Health Research Institute of Taiwan) and the Institutional Review Board of the Los Angeles Biomedical Research Institute. All participants provided written informed consent.

### Exome Sequencing, Variant calling, and QC

DNA samples from 6,809 SWAI individuals were exome sequenced at the Regeneron Genetics Center, using sequencing methodology, genome alignment, and genotype calling approaches as previously describe^36^. Briefly, exonic regions were targeted using an xGEN probe library with slight modifications. Targeted DNA was sequenced on the Illumina HiSeq 2500 platform with v4 chemistry using 75bp paired-end reads. Sequencing was performed such that >85% of the bases were covered at ≥20x depth. Read alignment to human genome reference GRCh38 and variant calling were performed using BWA-MEM and GATK, respectively. 93 samples were removed based on QC metrics including low coverage (<75% of targeted bases with at least 20x depth), low quality, sex mismatch, sample duplicates, and high discordance with array genotypes, resulting in the final count of 6,716 exomes for analysis. Variants were further filtered by missing call rates (<10%) and Hardy-Weinberg equilibrium p-values (>1×10^−15^).

DNA samples from 29,575 individuals of European ancestry from the DiscovEHR study were exome sequenced by the same method. DNA samples from 13,947 individuals of East Asian ancestry from the TAICHI study were exome sequenced by an analogous method as previously described^34^, with the major difference being the use of VCRome reagent for exome targeting instead of xGEN reagent.

### Variant annotation

Variants were annotated for their predicted effects on all protein-coding transcripts with annotated start and stop in Ensembl85 (54,214 transcripts corresponding to 19,467 genes) using snpEff^37^. Variants were annotated as predicted loss-of-function (pLOF) when they were predicted to incur frameshift, premature stop codon, loss of start or stop codon, or disruption of canonical splice dinucleotides. Nonsynonymous variants included missense SNVs and inframe indels. When a variant had different predicted effects among different transcripts, a more deleterious effect was prioritized. The variants detected in the American Indian exomes were compared to dbSNP (v151)^38^ and gnomAD exomes (r2.1)^39^.

### Principal Component Analysis

Reference genomes were downloaded from 1000 Genomes project server^40^. The analysis was limited to autosomal biallelic variants with MAF ≥ 5% and r^2^ < 0.2 outside of the major histocompatibility complex region that were detected in both the reference genomes and the SWAI exomes. We first calculated the principal components from the reference genomes and projected individuals from the SWAI study on to the PC space using PLINK2^41^.

### Comparison of allelic architecture

The allelic architecture of SWAI exomes was compared to European ancestry exomes from the DiscovEHR study and East Asian exomes, predominantly of Han Chinese from Taiwan, from the TAICHI study. All studies were exome sequenced at the Regeneron Genetics Center, but two different exome targeting reagents were used (xGEN and VCRome). To account for the difference in the exome targeting reagents, all comparisons were made among the subset of variants that map to the intersection of consistently covered regions of each targeting reagent. Consistently covered regions are defined as having ≥20x read depth in ≥90% of a randomly sampled set of 1,000 exomes sequenced using the targeting reagent.

For the comparison of proportional site frequency spectra with 6,716 European exomes, 6,716 East Asian exomes were randomly extracted from DiscovEHR and TAICHI studies, respectively. The number of pLOF and nonsynonymous variants were counted according to the minor allele count bins and the proportion was calculated.

For comparisons of allele frequency, we included only self-reported full American Indians from the SWAI study, to minimize the impact of admixture. To avoid situations where the minor allele of the same variant differs between studies, all allele frequencies refer to the alternate allele frequencies (AAF) of the variant compared to the human genome reference. For any study, if no alternate alleles were observed within a consistently covered region (as described above), the allele frequency of the variant in that study was inferred to be 0. SWAI allele frequencies were also compared to the population frequencies from gnomAD exomes r2.1. When a variant was not listed in gnomAD exomes, but the genomic position was called with mean read depth ≥20, the allele frequency of the variant in gnomAD was inferred to be 0.

### Association Tests

We derived the set of GWAS candidate effector genes from previous GWAS studies for body mass index^42^, type 2 diabetes^43^, and plasma lipid levels^44^. Sentinel variants of independent association signals were derived using GCTA-COJO^45^ using individuals of European ancestry from DiscovEHR as reference. The genes that are closest to the variants were derived using BEDTools and tested for association with corresponding traits in the American Indians.

For gene-burden tests, pLOF and missense variants were grouped into eight masks using two allele frequency cutoffs (AAF <1% and <5%) and four functional effect criteria: 1) M1 - pLOF variants only, 2) M2 - pLOF and all missense variants, 3) M3 - pLOF and missense variants predicted to be deleterious by all five prediction algorithms used (SIFT^46^, LRT^47^, MutationTaster^48^, PolyPhen2-HumDiv, PolyPhen2-HumVar^49^), 4) M4 - pLOF and missense variants predicted to be deleterious by at least one of the five prediction algorithms. If different masks of a gene are comprised of the same variants, they are collapsed to one mask with most stringent definition, so that only unique masks were tested for association. The Bonferroni corrected P-values were calculated as 0.05 / total number of unique masks tested. For masks with significant associations, the individual variants that were included in those masks were also tested for associations. Only the masks and variants with at least 10 alternate allele counts were tested.

Associations were tested under linear mixed model using SAIGE^50^ for diabetes status and BOLT^51^ for quantitative traits to adjust for population structure and cryptic relatedness. For diabetes, age, age^2^, sex, and 5 principal components of ancestry were included as covariates. For age of diabetes onset, sex and 5 principal components were included as covariates. Triglyceride measures were natural log transformed. For BMI and lipid traits, residuals were derived adjusting for age, age^2^, sex, 5 principal components, and transformed to normality by rank-based inverse normal transformation.

## Results

### Characterization of exome variants

We detected a total of 1,208,812 variants from the exomes of 6,716 SWAI (Table 1, Figure 1A), of which 1,130,961 (93.6%) were single nucleotide variants (SNVs) and 77,851 (6.4%) were indels. When annotated for predicted effects, 16,880 (1.4%) were predicted loss-of-function (pLOF) variants (frameshift, stop-gain, start-loss, splice acceptor, splice donor, and stop-loss) and 258,306 (21.4%) were nonsynonymous variants (inframe indels and missense). The majority of variants were rare, i.e., less than 10 alternate allele counts (corresponding to the alternate allele frequency of <0.07%) in SWAI.

**Table 1:** Summary statistics and annotation of variants captured by whole exome sequencing of 6,716 American Indians. Variants detected in SWAI exomes were categorized by their type and predicted functional effect. The number of variants were counted based on whether they have alternate allele count ≥10 in SWAI exomes and whether they are not present in dbSNP or gnomAD exomes.

| Variant type | | All | | | Alternate allele count $\geq 10$ | | |
| --- | --- | --- | --- | --- | --- | --- | --- |
|  |  | Total Number | Number (%) not in dbSNP<sup>a</sup> | Number (%) not in gnomAD<sup>b</sup> | Total Number | Number (%) not in dbSNP<sup>a</sup> | Number (%) not in gnomAD<sup>b</sup> |
| All |  | 1,208,812 | 245,039 (20.3%) | 545,979 (45.2%) | 393,548 | 76,966 (19.6%) | 175,318 (44.5%) |
| SNVs |  | 1,130,961 | 228,981 (20.2%) | 505,888 (44.7%) | 366,309 | 72,909 (19.9%) | 162,486 (44.4%) |
| Indels |  | 77,851 | 16,058 (20.6%) | 40,091 (51.5%) | 27,239 | 4,057 (14.9%) | 12,832 (47.1%) |
| Variant effect | | All | | | Alternate allele count $\geq 10$ | | |
|  |  | Total Number | Number (%) not in dbSNP<sup>a</sup> | Number (%) not in gnomAD<sup>b</sup> | Total Number | Number (%) not in dbSNP<sup>a</sup> | Number (%) not in gnomAD<sup>b</sup> |
| pLOF (N=16,880) | Frameshift | 6,881 | 2,456 (35.7%) | 2,552 (37.1%) | 1,474 | 401 (27.2%) | 418 (28.4%) |
|  | Stop gained | 5,288 | 1,427 (27.0%) | 1,659 (31.4%) | 1,016 | 315 (31.0%) | 354 (34.8%) |
|  | Start lost | 668 | 125 (18.7%) | 159 (23.8%) | 177 | 33 (18.6%) | 43 (24.3%) |
|  | Splice acceptor | 1,858 | 675 (36.3%) | 750 (40.4%) | 596 | 185 (31.0%) | 198 (33.2%) |
|  | Splice donor | 1,858 | 612 (32.9%) | 741 (39.9%) | 465 | 175 (37.6%) | 209 (44.9%) |
|  | Stop lost | 327 | 117 (35.8%) | 123 (37.6%) | 123 | 59 (48.0%) | 61 (49.6%) |
| Nonsynonymous<br>(N=258,306) | Inframe<br>indel | 4,157 | 591<br>(14.2%) | 801<br>(19.3%) | 1,323 | 119<br>(9.0%) | 173<br>(13.1%) |
|  | Missense | 254,149 | 40,529<br>(15.9%) | 49,061<br>(19.3%) | 68,494 | 11,088<br>(16.2%) | 12,631<br>(18.4%) |
| Synonymous |  | 164,772 | 16,650<br>(10.1%) | 20,898<br>(12.7%) | 54,952 | 4,551<br>(8.3%) | 5,357<br>(9.7%) |

<sup>a</sup> dbSNP v151 was used for comparison.
<sup>b</sup> gnomAD exomes r2.1 was used for comparison.

**Figure 1:**
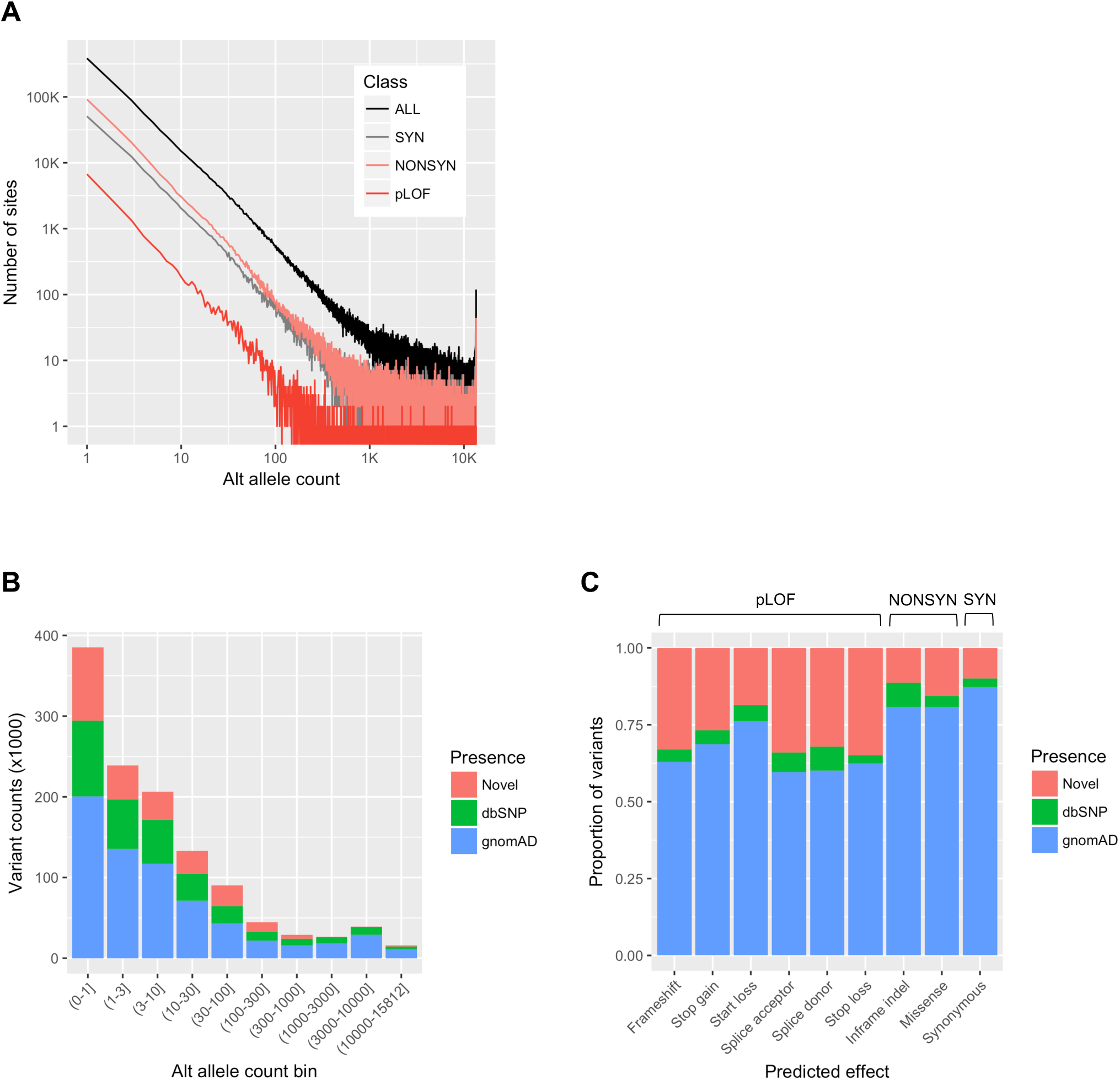
Summary statistics and annotation of variants captured by whole exome sequencing of 6,716 American Indians. (A) Site frequency distribution of 1,208,812 autosomal variants according to predicted functional effects. (B) The number of variants that are novel or previously listed in gnomAD or dbSNP databases as a function of alternate allele count. (C) The proportion of variants that are novel or previously listed in gnomAD or dbSNP databases stratified by predicted functional effect. pLOF: predicted loss-of-function, NONSYN: nonsynonymous, SYN: synonymous.

When compared to dbSNP and gnomAD exome databases, 241,042 variants (19.9%) were novel and were not listed in either database (20.3% not in dbSNP and 45.2% not in gnomAD exome). The novel variants tended to be rarer (Figure 1B) and more enriched among pLOF variants than among nonsynonymous or synonymous variants (Figure 1C).

### Population structure

SWAI study population has considerable admixture according to the self-reported American Indian ancestry of the study subjects: 72.8% of the subjects were full American Indians (all eight great grandparents were American Indian) while the rest had varying degrees of admixture (Figure S1A). To evaluate the population structure and admixture of SWAI based on the genetic data, we constructed principal components from three ancestral super populations (EUR, EAS, AFR) from the 1000 Genomes Project and projected SWAI study subjects onto the principal component space. When only the self-reported full American Indians were plotted, they clustered about an axis between European and East Asian clusters (Figure S1B). When all individuals from SWAI study were plotted, we observed that individuals with greater self-reported admixture tended to deviate further from the full American Indian cluster towards European and African clusters (Figure S1C). These results are consistent with study population being comprised of individuals with complete or partial American Indian ancestry.

### Comparison of allelic architecture and frequency

We compared the allelic architecture of SWAI exomes to European ancestry exomes from the DiscovEHR study and East Asian exomes from the TAICHI study that served as the extant proxies for ancestral European and East Asian genomes that influenced the American Indian genome. To account for the difference in exome targeting reagents across the studies (SWAI and DiscovEHR studies with xGEN and TAICHI study with VCRome), analyses were restricted to variants that reside in the consistently covered regions by both targeting reagents. We compared the proportional site frequency spectra of SWAI exomes to the same number of European and East Asian ancestry exomes that were randomly sampled. SWAI exomes were relatively depleted of ultra-rare pLOF and nonsynonymous variants (MAC ≤3) compared to European ancestry exomes but were enriched for moderately rare pLOF and nonsynonymous variants (3< MAC ≤1000) compared to both European and East Asian ancestry exomes (Figure 2A and 2B).

**Figure 2:**
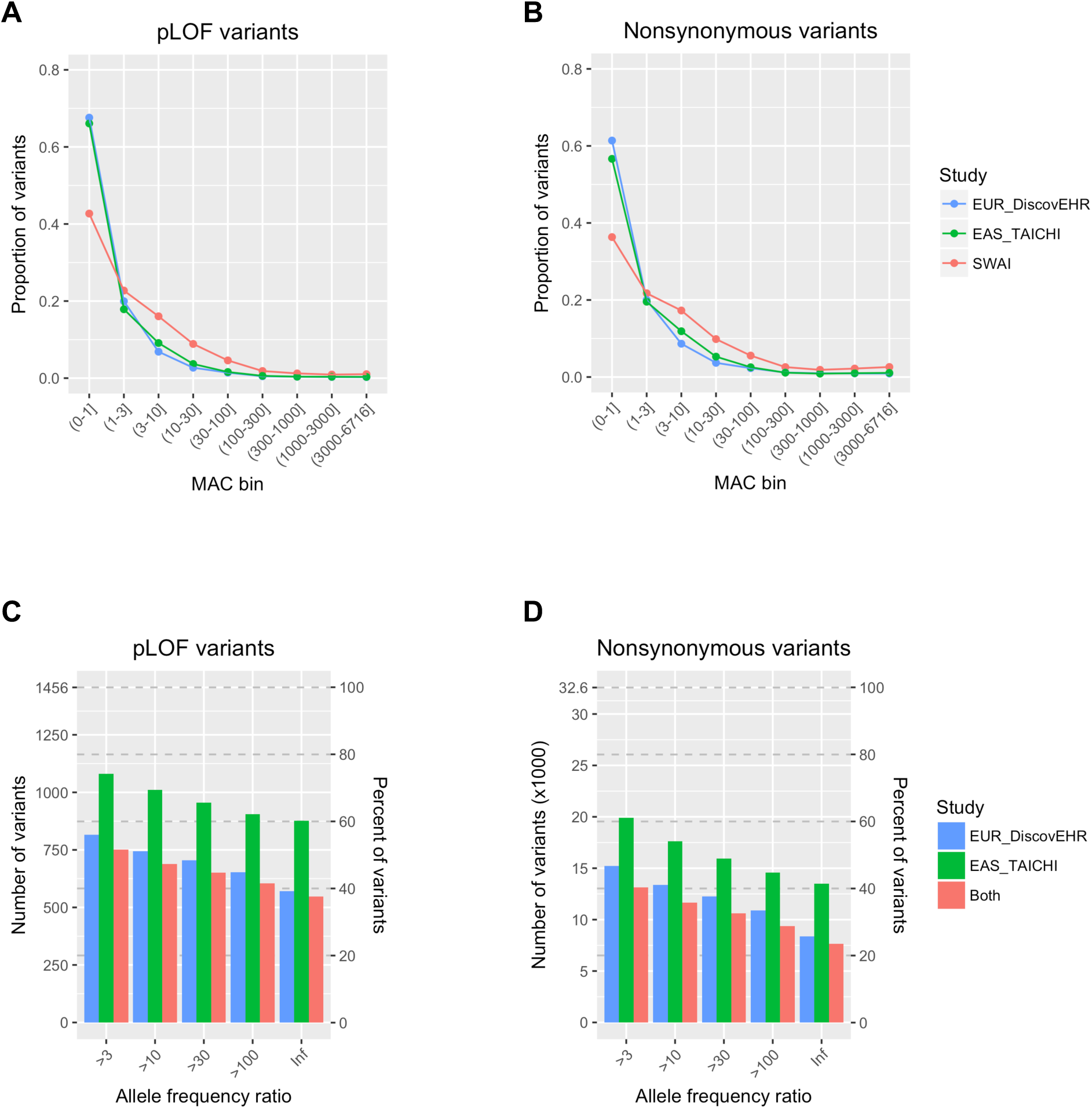
Comparison of the distribution and frequency of pLOF and nonsynonymous variants among SWAI, European, and East Asian exomes. (A-B) Comparison of the distribution of pLOF (A) and nonsynonymous (B) variants at different minor allele count (MAC) bins among American Indian, European, and East Asian exomes from SWAI, DiscovEHR, and TAICHI studies, respectively. (C-D) The number and percentage of pLOF (C) and nonsynonymous (D) variants that are enriched in full American Indian exomes from SWAI compared to European or East Asian exomes, or both European and East Asian exomes. The analysis is restricted to variants with alternate allele count ≥10 in full American Indians from SWAI. All analyses were restricted to variants in consistently covered regions to account for the difference in exome targeting reagents among the studies.

To examine how many of the variants that were detected in SWAI are private or enriched in SWAI, we compared the allele frequency of pLOF and nonsynonymous variants in full American Indians from SWAI to individuals with European and East Asian ancestries. The analysis was restricted to variants with minimum alternate allele count of 10 in SWAI, considering the power for statistical inference, within the consistently covered regions. Among the total of 1,456 pLOF variants, 548 (38.4%) were only detected in American Indians and 689 (48.3%) were more than 10 times more enriched in American Indians compared to both the individuals of European ancestry and the individuals of East Asian ancestries (Figure 2C). Among the total of 32,577 nonsynonymous variants, 7,640 (23.7%) were only detected in American Indians and 11,649 (36.1%) were more than 10 times more enriched in American Indians compared to individuals with European and East Asian ancestries (Figure 2D).

### Genes with pLOF variation

As predicted loss-of-function variants can provide a valuable insight on the biological connection between genes and traits, we examined how many genes carried pLOF variation in SWAI exomes. Of the 19,467 genes annotated, 9,015 genes (46.3%) had at least one heterozygous carrier of pLOF variants and 3,398 genes (17.5%) had at least 10 heterozygous carriers (Table 2, Figure 2A). 907 genes (4.7%) had at least one homozygous carrier of pLOF variants, and 466 genes (2.4%) had at least 10 homozygous carriers.

**Table 2:** Number and percentage of genes with at least X number of carriers of predicted loss-of-function variants in 6,716 SWAI exomes.

| **Number of carriers** | **Number (%) of genes with any carriers** | **Number (%) of genes with heterozygous carriers** | **Number (%) of genes with homozygous carriers** |
| --- | --- | --- | --- |
| ≥1 | 9,016 (46.3%) | 9,015 (46.3%) | 907 (4.7%) |
| ≥3 | 5,910 (30.4%) | 5,907 (30.3%) | 593 (3.0%) |
| ≥10 | 3,407 (17.5%) | 3,398 (17.5%) | 466 (2.4%) |
| ≥30 | 1,948 (10.0%) | 1,936 (9.9%) | 389 (2.0%) |
| ≥100 | 953 (4.9%) | 939 (4.8%) | 327 (1.7%) |

To see whether population history impacted the number and distribution of pLOF variation, we compared the number of genes with pLOF carriers in the same number of samples randomly drawn from American Indian, European, and East Asian exomes from SWAI, DiscovEHR and TAICHI studies. The analysis was again restricted to variants in the consistently covered regions across the studies for comparison. Consistent with founder effect, the number of genes with heterozygous pLOF carriers was lower in SWAI exomes than in European and East Asian exomes (Figure 2B, top). On the other hand, the number of genes with homozygous pLOF carriers was greater in SWAI exomes (Figure 2B, bottom), potentially due to the fact that SWAI population underwent reproductive isolation with small population size.

pLOF variation may accumulate due to random genetic drift or specific environmental pressure that populations face which could increase tolerance to loss-of-function of certain genes. We investigated the overlap among the set of genes with ≥10 pLOF carriers in the SWAI (N = 6,716), European (N = 29,575) and East Asian exomes (N = 13,947). We set the minimum number of carriers at 10 considering the power for downstream statistical inference. While the total sample size of SWAI exomes was smaller than those of European and East Asian exomes, there were 275 genes with ≥10 heterozygous pLOF carriers and 87 genes with ≥10 homozygous carriers only in SWAI exomes and not others (Figure S2). Of all the genes with ≥10 heterozygous and ≥10 homozygous pLOF carriers in SWAI exomes, ∼11.8% and 27.7% were unique to SWAI exomes, respectively.

### Association with metabolic traits

Genetic associations in American Indians using exome variants can not only provide additional evidence for the candidate effector genes in GWAS loci, but also find novel variants, with potential clinical impact, that are unique or enriched in American Indians. We derived the list of candidate effector genes from the latest and largest GWAS studies for body mass index (BMI), type 2 diabetes, and plasma lipid levels and tested their association with respective traits in American Indians. We used gene-burden approach, aggregating pLOF and nonsynonymous variants into eight masks, using two allele frequency cutoffs (<1% and <5%, indicated as 1 and 5 following ‘.’ in the name of the mask) and four functional effect criteria: 1) M1 - pLOF variants only, 2) M2 - pLOF and all missense variants, 3) M3 - pLOF and missense variants predicted to be deleterious by all five prediction algorithms used (see methods for detail), 4) M4 - pLOF and missense variants predicted to be deleterious by at least one of the five prediction algorithms. If different masks of a gene are comprised of the same variants, they are collapsed to one mask with most stringent definition, so that only unique masks were tested for association. The Bonferroni corrected P-values were calculated by dividing 0.05 by the number of unique masks tested.

## Body mass index

774 genes that were closest to the independent association signals in the latest BMI GWAS study^42^ were analyzed for association with maximum BMI measured in American Indians (Bonferroni P < 0.05 / 1922 unique masks = 2.6 ×10^−5^). The M3.1 mask of *MC4R* [OMIM: 155541], a known gene for early-onset obesity, was the only one significantly associated with increased maximum BMI in SWAI (Table 3, Beta = 0.56sd, P = 5.2×10^−9^). The M3 mask consisted of seven variants, including the previously described frameshift variant (p.Gly34fs), and missense variants (p.Arg165Gly, p.Ala303Pro, and p.Arg165Gln) that are either private or enriched in American Indians and were associated with maximum BMI individually (Table S1)^27^. These variants were previously identified by targeted sequencing of *MC4R* in SWAI and were found to impair the activity of MC4R *in vitro*, suggesting their functional impact^27^.

**Table 3:** Gene-burden associations of candidate GWAS effector genes for body mass index, type2 diabetes, and plasma lipid levels with respective traits in American Indians.

|  | Association results from GWAS<sup>42</sup> |  |  |  |  | Gene-level associations in SWAI |  |  |  |  |
| --- | --- | --- | --- | --- | --- | --- | --- | --- | --- | --- |
| Trait | Body mass index |  |  |  |  | Maximum BMI |  |  |  |  |
| Gene | Top variant | Variant effect | AAF<sup>a</sup> | Beta<sup>b</sup> | P value | Top mask<sup>d</sup> | Freq | Beta<sup>b</sup> | P value |  |
| MC4R | rs6567160 | intergenic | 0.23 | 0.055 | 1.8E-178 | M3.1 | 0.011 | 0.562 | 5.2E-09 |  |
| Trait | Type 2 diabetes |  |  |  |  |  |  |  |  |  |
| Gene | Top variant | Variant effect | AAF | OR<sup>c</sup> | P value | Top mask | Freq | OR<sup>c</sup> | P value |  |
| MC4R | rs523288 | intergenic | 0.238 | 1.05 | 7.6E-13 | M3.1 | 0.010 | 2.62 | 1.2E-05 |  |
| ABCC8 | rs67254669 | missense | 0.001 | 1.89 | 1.1E-08 | M3.5 | 0.018 | 2.21 | 9.3E-06 |  |
| Trait | Plasma lipid levels |  |  |  |  |  |  |  |  |  |
| Gene | Top variant | Variant effect | AAF | Trait | Beta | P value | Top mask | Freq | Beta | P value |
| APOB | rs541041 | intergenic | 0.81 | TC | 0.11 | 5.3E-237 | M4.5 | 0.062 | -0.208 | 1.4E-06 |
|  |  |  |  | LDLC | 0.12 | 1.3E-287 |  |  | -0.286 | 1.6E-09 |
| APOE | rs445925 | downstream | 0.11 | TC | -0.21 | 0 | M4.1 | 0.014 | -0.630 | 9.4E-14 |
|  |  |  |  | LDLC | -0.32 | 0 |  |  | -0.860 | 4.7E-20 |
| PCSK9 | rs11591147 | missense | 0.015 | TC | -0.41 | 0 | M2.5 | 0.057 | -0.202 | 6.2E-06 |
|  |  |  |  | LDLC | -0.48 | 0 | M3.5 | 0.028 | -0.443 | 9.1E-11 |
| TM6SF2 | rs58542926 | missense | 0.074 | TC | -0.13 | 7.0E-155 | M4.5 | 0.067 | -0.256 | 4.6E-10 |
|  |  |  |  | LDLC | -0.10 | 6.5E-93 |  |  | -0.223 | 7.9E-07 |
|  |  |  |  | TG | -0.12 | 3.7E-125 |  |  | -0.283 | 2.5E-11 |
| GPAM | rs2792751 | missense | 0.73 | HDLC | -0.03 | 3.8E-21 | M3.5 | 0.026 | 0.583 | 5.1E-15 |
| *LIPC* | rs1800588 | upstream | 0.24 | HDLC | 0.12 | 0 | M4.1 | 0.013 | 0.466 | 9.8E-6 |
| *LIPG* | rs7241918 | intergenic | 0.85 | HDLC | 0.08 | 1.2E-104 | M3.1 | 0.002 | 1.279 | 2.7E-5 |

<sup>a</sup> AAF: alternate allele frequency
<sup>b</sup> The unit is standard deviation of normalized traits.
<sup>c</sup> OR: odds ratio
<sup>d</sup> The mask with strongest trait association is displayed. Refer to the methods for detailed mask definition.

## Type 2 diabetes

269 genes that were closest to the independent association signals in the latest T2D GWAS study^43^ were analyzed for association with T2D in American Indians (Bonferroni P < 0.05 / 772 unique masks = 6.5×10^−5^). Two masks were significantly associated with T2D risk: M3.1 mask of *MC4R* and M3.5 mask of *ABCC8* [OMIM: 600509], a known gene for maturity onset diabetes of young (MODY) [OMIM: 606391] (Table 3).

The same M3.1 mask of *MC4R*, which was associated with maximum BMI, was also associated with T2D (OR = 2.6, P = 1.2×10^−5^). When adjusted for maximum BMI, the association was only partially mitigated (OR = 2.2, P = 5.8×10^−4^), suggesting that *MC4R* may affect T2D independent of its effect on obesity. Again, individual variants in the mask that are unique or highly enriched in American Indians, p.Gly34fs, p.Arg165Gly, and p.Arg165Gln, were associated with increased T2D risk (Table S2). The mask was also associated with earlier onset of T2D (Beta = −4.3years, P = 5.5×10^−3^), with all three homozygous carriers developing T2D under the age of 30 years (Figure S3A).

The M3.5 mask of *ABCC8* was associated with diabetes (OR = 2.2, P = 9.3×10^−6^). Among the 17 variants included in the M3.5 mask of *ABCC8*, p.Arg1420His was most strongly associated with diabetes risk (OR = 2.2, P = 1.5×10^−5^), which was previously reported^30^. Notably, this variant was ∼489-fold and ∼115-fold enriched in full American Indians compared to individuals with European ancestry and those with East Asian ancestry, respectively (Table S2). Consistent with the known role of *ABCC8* in MODY, early-onset form of diabetes, and what was previously reported for p.Arg1420His alone, the M3.5 mask was associated with earlier age of onset (Beta = −6.9years, P = 1.8×10^−7^), with the one homozygous carrier developing diabetes before the age of 10 (Figure S3B). *ABCC8* encodes sulfonylurea receptor 1 protein (SUR1) that constitutes ATP-sensitive potassium (K_ATP_) channel and it was previously shown that the p.Arg1420His mutation in SUR1 protein leads to impaired activity of K_ATP_ channel *in vitro*^30^, suggesting the functional impact of the variant.

## Plasma lipids

Up to 115 genes that were closest to the independent association signals in the latest GWAS study for plasma lipid traits^44^ were analyzed for association with fasting total cholesterol, HDL cholesterol, LDL cholesterol, and triglyceride levels in American Indians (Bonferroni P < 0.05 / up to 391 unique masks = 1.28 ×10^−4^). Seven genes were significantly associated with at least one lipid trait (Table 3), among which six genes, *APOB, APOE, PCSK9, TM6SF2, LIPC*, and *LIPG*, have well characterized roles in lipid metabolism. On the other hand, *GPAM* gene, which encodes mitochondrial glycerol-3-phosphate acyltransferase with no previously known role in HDL metabolism, was associated with increased HDL cholesterol levels under M3.5 mask (Beta = 0.58sd, P = 5.1×10^−15^). Among 15 variants included in the M3.5 mask, p.Ser611Arg variant was most strongly associated with HDL cholesterol levels (Beta = 0.57sd, P = 3.8×10^−14^). The p.Ser611Arg variant is present in American Indians at AAF of 0.025, but not detected in individuals with European ancestry and ∼383-fold enriched compared to individuals with East Asian ancestry (Table S3). This finding complements the previously identified association of a common missense variant of *GPAM*, p.Ile43Val, with HDL cholesterol levels in European populations^44^.

Notably, the *APOB, APOE, PCSK9*, and *TM6SF2* masks that were associated with LDL cholesterol levels included many variants that are enriched or private in SWAI and have large effect sizes on LDL cholesterol levels (Table S3), suggesting clinical impacts of these variants. A frameshift pLOF variant of *APOB*, p.Ala3175fs, is private in SWAI (AAF = 0.001) and was associated with lower LDL cholesterol levels (Beta = −2.30sd, P = 1.8×10^−13^). A missense variant of *APOE*, p.Ala184Asp, is private in SWAI (AAF = 0.007) and was associated with lower LDL cholesterol levels (Beta = −1.18sd, P = 2.3×10^−20^). This variant was in linkage equilibrium with the common variants of *APOE* e2 and e4 haplotypes (r2 < 0.05). A missense variant of *PCSK9*, p.Gly244Asp, is highly enriched in SWAI (AAF = 0.024) and was associated with lower LDL cholesterol levels (Beta = − 0.46sd, P=4.7×10^−10^). A missense variant of *TM6SF2*, p.Arg138Trp, is highly enriched in SWAI (AAF = 0.046) and was associated with lower LDL cholesterol levels (Beta = − 0.20sd, P=1.2×10^−4^). Further studies are needed to demonstrate the functional impacts of these variants and evaluate their implications in cardiovascular health.

## Discussion

Our study illustrates that exome sequencing applied to founder populations such as American Indians can uncover novel genetic variations that are associated with clinical and quantitative traits and expand our understanding of the genetic contribution to these traits. This is enabled by the distinct allelic architecture of American Indians with rare functional variants drifted to higher frequency, increasing the statistical power to detect their associations with traits. In addition, gene-burden approaches, aggregating rare pLOF and nonsynonymous variants affecting the same gene, further enhanced the power to evaluate the relationship between genes and traits of interest.

The genetic architecture of the SWAI is influenced by their unique population history involving bottleneck events followed by isolation. Consistent with the expectation that bottleneck events reduce overall genetic diversity, we observed fewer numbers of pLOF and nonsynonymous variants in SWAI exomes compared to European and East Asian exomes that underwent rapid population growth. Isolation subsequent to bottleneck events can randomly increase the frequency of rare variants. When we compared the proportion of pLOF and nonsynonymous variants across minor allele count bins, we observed selective enrichment of moderately rare variants in American Indian exomes compared to European and East Asian ancestry exomes, similar to the observation in Finnish populations that also underwent a series of bottleneck events and isolation^5^. In addition, reproductive isolation in small populations can increase the homozygosity of genetic variants. As expected, SWAI had greater number of pLOF and nonsynonymous variants in homozygosis compared to equivalent numbers of more cosmopolitan European and East Asian ancestry populations.

Genome-wide association studies have traditionally focused on common variants that are captured by genotyping arrays or imputation and, as a result, many association signals are noncoding, making it challenging to pinpoint the effector genes that mediate the association. In our study, we took the candidate genes from large GWAS studies conducted for BMI, type 2 diabetes, and plasma lipid traits in European populations and tested their association in American Indians using gene-burden approach. We found significant associations for a handful of these genes, providing additional evidence for the connection between these candidate genes and the traits. Of note, gene-burden associations tended to have stronger effects on traits compared to associations found in GWAS, consistent with the expectation that rare pLOF and nonsynonymous variants have greater impacts than common noncoding variants (Table 3). Most of the associations were with genes that have well characterized roles in the regulation of the respective traits. One exception was the association of *GPAM* gene, which encodes mitochondrial glycerol-3-phosphate acyltransferase, with plasma HDL-cholesterol levels, implicating a potential novel role of *GPAM* in HDL metabolism. Notably, a previous study on *Gpam* knockout mice observed reduced hepatic triglyceride content and plasma total cholesterol and triglyceride levels, but no significant difference in plasma HDL cholesterol levels^52^, suggesting that the effect of *GPAM* on HDL cholesterol may be specific to humans.

The current study using the whole exome sequence of SWAI complements and extends previous genetic studies that have been conducted in SWAI using targeted sequencing or genotyping of candidate genes and variants and high-density genotyping arrays. The whole-exome sequence enabled the systemic examination of all candidate genes for their association with metabolic traits at the gene-level, which confirmed significant associations of *MC4R* and *ABCC8* for BMI and T2D that were previously found in SWAI by targeted sequencing of these specific genes^27; 30^. In addition, the whole exome sequence allowed the identification of rare coding variants beyond the common variants that have been captured by targeted genotyping or genotyping arrays^22; 23; 26^, leading to a more comprehensive understanding of the impact of genetic variation in the candidate genes on traits. A previous GWAS study for T2D performed in SWAI using a genotyping array found genome-wide significant associations of two common intronic variants in *KCNQ1* and *DNER* with T2D risk^17; 21^. We did not find additional association of pLOF or nonsynonymous variants of *KCNQ1* and *DNER* with T2D risk, suggesting that the previously observed GWAS association signals are likely mediated by alteration in transcriptional regulation.

It is worth noting that most gene-burden associations that we found were driven by pLOF and/or nonsynonymous variants that are unique or highly enriched in American Indians. Many of these variants were associated with traits with strong effects, warranting further investigation on the clinical implications of these variants in American Indians. In addition, further characterization of the functional impact of these protein-sequence altering variants can broaden our understanding of the structure and regulation of the proteins. While the current study specifically focused on the exome variants within the GWAS candidate genes, more studies are ongoing to identify novel genetic associations utilizing exome variants across the genome and could shed light on additional genetic underpinnings of the high prevalence of metabolic disorders in this population.

## Supplemental Data Description

Supplemental data include 3 figures and 3 tables.

**Figure 3:**
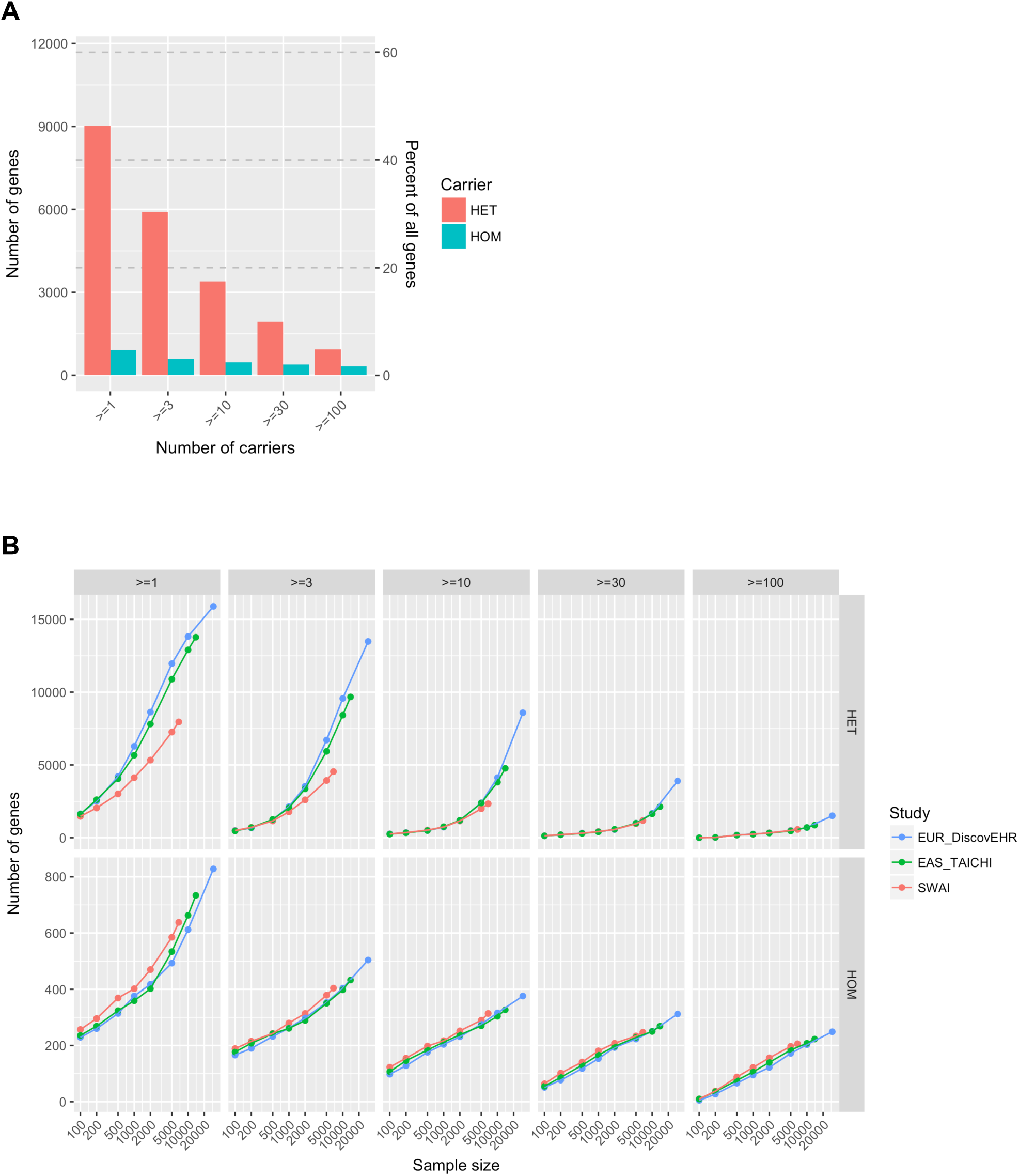
Comparison of the number of genes with predicted loss-of-function carriers among SWAI, European, and East Asian exomes. (A) The number and percentage of genes among 19,467 annotated autosomal genes with at least X number of heterozygous and homozygous pLOF carriers in SWAI study alone. (B) The comparison of the number of genes with at least X number of heterozygous (top) and homozygous (bottom) pLOF carriers at fixed sample sizes randomly extracted from SWAI, European, and East Asian exomes. The analysis was restricted to the variants in consistently covered regions to account for the difference in exome targeting reagents among the studies.

## Declaration of Interests

H.K., N.G., B.Y., S.K., A.R.S., C.V.H. are current or former employees and/or stock holders of Regeneron Genetics Center or Regeneron Pharmaceuticals. The other authors declare no competing interests.

## Acknowledgements

We thank the volunteers from the American Indian community who participated in the study, the participants of the DiscovEHR study, and the participants of the TAICHI study. We also thank all research staff and teams at the Regeneron Genetics Center, National Institute of Diabetes and Digestive and Kidney Diseases, Geisinger Health System, and TAICHI consortium, who contributed to the current study. The study is funded by Regeneron Pharmaceuticals and the Intramural Research Program of the National Institute of Diabetes and Digestive and Kidney Diseases.

## Web Resources

dbSNP, https://www.ncbi.nlm.nih.gov/snp

gnomAD, https://gnomad.broadinstitute.org

PLINK2, www.cog-genomics.org/plink/2.0

1000 Genomes Projects, https://www.internationalgenome.org

OMIM, http://www.omim.org

