## Supplemental data for "Characterization of exome variants and their metabolic impact in 6,716 American Indians from Southwest US"

### Figure S1: Population structure in the SWAI study

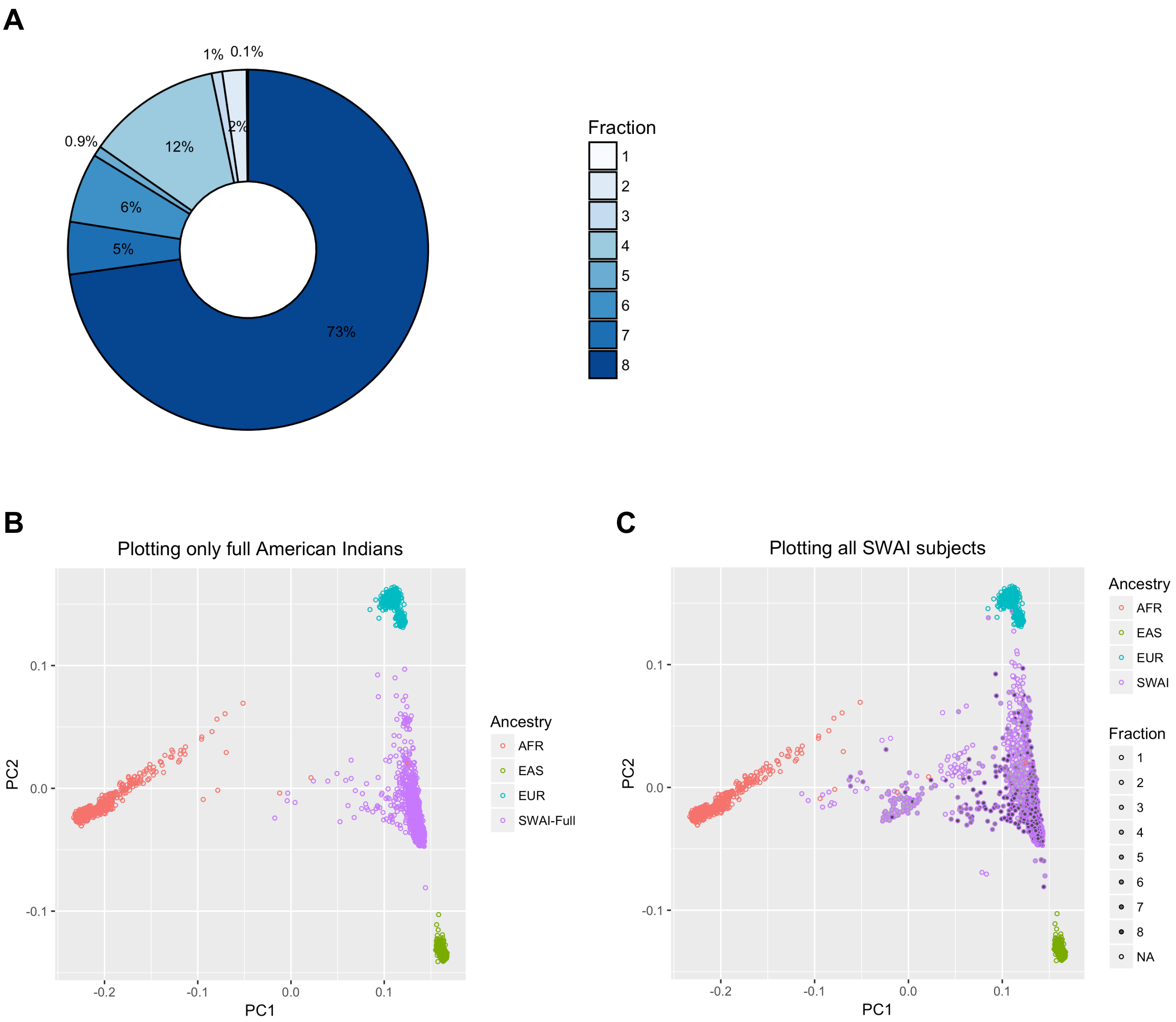


(A) The percentage of the individuals according to the self-reported admixture (fraction of great grandparents that were American Indian out of 8). (B) Individuals with full self-reported American Indian ancestry in the SWAI study were projected onto the PC space calculated from African, East Asian, and European ancestries from 1000 genomes project. (C) All individuals from the SWAI study were projected onto the PC space with the fill intensity indicating their self-reported admixture.

**Figure S2: Overlap of genes with at least 10 heterozygous or homozygous pLOF carriers among SWAI, European, and East Asian exomes**

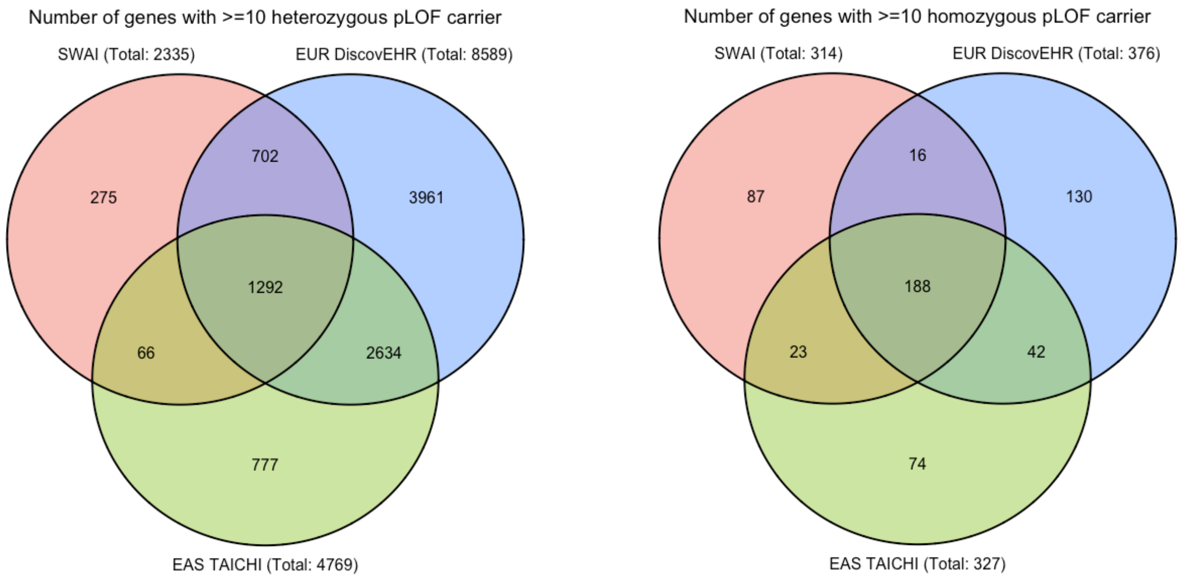


The Venn diagrams represent the overlap of the genes with at least 10 heterozygous (left) and homozygous (right) pLOF carriers in 6,716 SWAI, 29,575 European, and 13,947 East Asian exomes.

### Figure S3: The distribution of age of onset per genotype

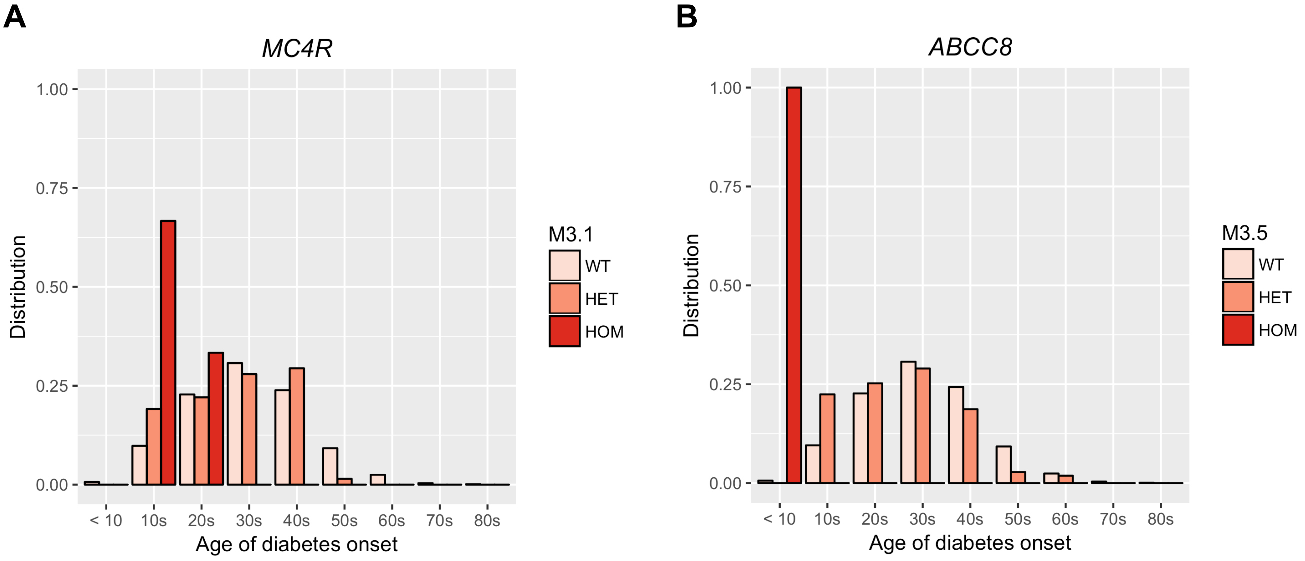


The distribution of age of diabetes onset per *MC4R* M3.1 mask genotype (A) and *ABCC8* M3.5 mask genotype (B).

### Table S1: Association of individual variants within the *MC4R* M3.1 mask with maximum body mass index

| Gene Mask | Variant[a] | Variant effect | Amino acid change | Beta[b] | P value | AAF[c] | Allele frequency ratio (AAF in full American Indians / AAF in respective populations) | | | | | | |
|---|---|---|---|---|---|---|---|---|---|---|---|---|---|
| | | | | | | | GHS EUR | TAICHI EAS | gnomAD ALL | gnomAD AMR | gnomAD EUR | gnomAD EAS | gnomAD AFR |
| *MC4R* M3.1 | 18:60371857:G:C | missense | p.Arg165Gly | 0.666 | 9.8E-04 | 0.0025 | Inf | NC | Inf | Inf | Inf | Inf | Inf |
| | 18:60372250:C:CT | frameshift | p.Gly34fs | 0.622 | 1.0E-03 | 0.0023 | Inf | Inf | Inf | Inf | Inf | Inf | Inf |
| | 18:60371443:C:G | missense | p.Ala303Pro | 0.881 | 1.0E-02 | 0.0008 | Inf | Inf | Inf | Inf | Inf | Inf | Inf |
| | 18:60371856:C:T | missense | p.Arg165Gln | 0.353 | 2.0E-02 | 0.0044 | 132.9 | 125.3 | 188.2 | Inf | 102.1 | 82.6 | Inf |

[a] Variant is designated as chromosome:position:reference allele:alternate allele

[b] The unit is standard deviation of normalized traits.

[c] AAF: alternate allele frequency

The individual variants included in the *MC4R* M3.1 mask with ≥10 alternate allele counts were tested for association with maximum body mass index in SWAI. The allele frequency ratios were derived by dividing the allele frequency in the full American Indians from the SWAI study by the allele frequencies in the European and East Asian individuals from DiscovEHR and TAICHI studies, and ancestral populations from the gnomAD exome database. When variants were not captured and fall outside of the consistently covered region in a population, they were denoted as “not captures (NC)”.

**Table S2: Association of individual variants within the *MC4R* M3.1 and *ABCC8* M3.5 masks with diabetes**

| Gene Mask | Variant[a] | Variant effect | Amino acid change | OR[b] | P value | AAF[c] | Allele frequency ratio (AAF in full American Indians / AAF in respective populations) | | | | | | |
|---|---|---|---|---|---|---|---|---|---|---|---|---|---|
| | | | | | | | GHS EUR | TAICHI EAS | gnomAD ALL | gnomAD AMR | gnomAD EUR | gnomAD EAS | gnomAD AFR |
| *MC4R* M3.1 | 18:60372250:C:CT | frameshift | p.Gly34fs | 3.04 | 7.90E-03 | 0.0021 | Inf | Inf | Inf | Inf | Inf | Inf | Inf |
| | 18:60371857:G:C | missense | p.Arg165Gly | 2.92 | 1.61E-02 | 0.0024 | Inf | NC | Inf | Inf | Inf | Inf | Inf |
| | 18:60371856:C:T | missense | p.Arg165Gln | 2.08 | 3.91E-02 | 0.0043 | 132.9 | 125.3 | 188.2 | Inf | 102.1 | 82.6 | Inf |
| | 18:60371443:C:G | missense | p.Ala303Pro | 2.55 | 2.62E-01 | 0.0008 | Inf | Inf | Inf | Inf | Inf | Inf | Inf |
| *ABCC8* M3.5 | 11:17395658:C:T | missense | p.Arg1420His | 2.23 | 1.53E-05 | 0.0162 | 489.2 | 115.3 | 701.8 | 107.0 | Inf | Inf | Inf |

[a] Variant is designated as chromosome:position:reference allele:alternate allele

[b] OR: odds ratio

[c] AAF: alternate allele frequency

The individual variants included in the *MC4R* M3.1 and *ABCC8* M3.5 mask with ≥10 alternate allele counts were tested for association with diabetes in SWAI. The allele frequency ratios were derived by dividing the allele frequency in the full American Indians from the SWAI study by the allele frequencies in the European and East Asian individuals from DiscovEHR and TAICHI studies, and ancestral populations from the gnomAD exome database. When variants were not captured and fall outside of the consistently covered region in a population, they were denoted as “not captures (NC)”.

### Table S3: Association of individual variants within the masks of *APOB*, *APOE*, *PCSK9*, *TM6SF2*, *GPAM*, *LIPC*, and *LIPG* with plasma lipid levels

| Gene Mask | Variant[a] | Variant effect | Amino acid change | TC | | LDLC | | HDLC | | TG | | AAF[c] | Allele frequency ratio (AAF in full American Indians / AAF in respective populations) | | | | | | |
|---|---|---|---|---|---|---|---|---|---|---|---|---|---|---|---|---|---|---|---|
| | | | | Beta[b] | P value | Beta[b] | P value | Beta[b] | P value | Beta[b] | P value | | GHS EUR | TAICHI EAS | gnomAD ALL | gnomAD AMR | gnomAD EUR | gnomAD EAS | gnomAD AFR |
| *APOB* M4.5 | 2:21007344:GC:G | frameshift | p.Ala3175fs | -1.78 | 6.7E-10 | -2.30 | 1.8E-13 | 0.50 | 1.5E-01 | -1.03 | 4.2E-04 | 0.0012 | Inf | Inf | Inf | Inf | Inf | Inf | Inf |
| | 2:21006782:A:C | missense | p.Ile3362Met | -0.18 | 4.3E-03 | -0.27 | 8.7E-05 | 0.06 | 4.4E-01 | 0.10 | 1.2E-01 | 0.0262 | Inf | Inf | 2324.7 | 320.2 | Inf | Inf | Inf |
| | 2:21011100:T:C | missense | p.His1923Arg | -0.51 | 5.2E-03 | -0.66 | 1.1E-03 | -0.04 | 8.4E-01 | -0.12 | 5.1E-01 | 0.0031 | 0.0 | 4.3 | 0.0 | 0.1 | 0.0 | 4.2 | 0.1 |
| | 2:21026844:C:T | missense | p.Val730Ile | -0.28 | 6.5E-02 | -0.27 | 1.1E-01 | 0.18 | 3.2E-01 | -0.52 | 9.7E-04 | 0.0046 | 0.0 | 25.6 | 0.0 | 0.0 | 0.0 | Inf | 0.2 |
| | 2:21013525:C:T | missense | p.Arg1284Gln | -0.53 | 9.2E-02 | NA | NA | -0.85 | 2.1E-02 | NA | NA | 0.0010 | 78.5 | 37.0 | 111.1 | Inf | 150.7 | Inf | Inf |
| | 2:21037212:G:A | missense | p.Thr194Met | -0.31 | 1.8E-01 | -0.34 | 1.8E-01 | 0.23 | 4.1E-01 | -0.55 | 2.4E-02 | 0.0019 | 21.1 | 0.0 | 0.4 | 0.6 | 11.6 | 0.0 | 4.4 |
| | 2:21015541:C:G | missense | p.Asp1113His | -0.30 | 2.1E-01 | -0.40 | 1.2E-01 | -0.25 | 3.7E-01 | 0.17 | 4.9E-01 | 0.0018 | 0.0 | 4.3 | 0.0 | 0.1 | 0.0 | Inf | 0.2 |
| | 2:21006931:G:T | missense | p.Leu3313Ile | -0.24 | 3.0E-01 | -0.23 | 3.7E-01 | 0.21 | 4.4E-01 | -0.36 | 1.3E-01 | 0.0019 | Inf | Inf | 8.7 | 13.2 | Inf | Inf | 0.6 |
| | 2:21010635:T:C | missense | p.Gln2078Arg | -0.24 | 3.0E-01 | -0.23 | 3.7E-01 | 0.21 | 4.4E-01 | -0.36 | 1.3E-01 | 0.0019 | Inf | Inf | Inf | Inf | Inf | Inf | Inf |
| | 2:21011704:C:T | missense | p.Asp1722Asn | 0.07 | 5.7E-01 | 0.11 | 4.0E-01 | 0.03 | 8.3E-01 | -0.18 | 1.2E-01 | 0.0074 | 93.0 | 54.8 | 329.4 | 90.6 | 446.8 | Inf | Inf |
| | 2:21007773:G:T | missense | p.Thr3032Asn | -0.14 | 5.9E-01 | -0.37 | 1.8E-01 | 0.38 | 2.3E-01 | -0.18 | 5.0E-01 | 0.0015 | 36.2 | Inf | 30.7 | 10.6 | 69.5 | Inf | 9.9 |
| | 2:21009973:C:G | missense | p.Asp2299His | -0.14 | 6.8E-01 | -0.31 | 3.8E-01 | 0.25 | 5.2E-01 | 0.23 | 4.9E-01 | 0.0010 | 1.0 | Inf | 0.1 | 0.1 | 0.5 | Inf | 0.0 |
| | 2:21038062:G:A | missense | p.Pro145Ser | -0.08 | 7.3E-01 | -0.09 | 7.1E-01 | 0.06 | 8.2E-01 | -0.01 | 9.7E-01 | 0.0021 | 0.2 | Inf | 0.0 | 0.0 | 0.2 | Inf | 0.0 |
| | 2:21006988:A:G | missense | p.Ser3294Pro | -0.04 | 8.4E-01 | -0.20 | 4.1E-01 | 0.00 | 9.9E-01 | 0.05 | 8.2E-01 | 0.0022 | 1.7 | Inf | 0.1 | 0.1 | 1.0 | Inf | 0.0 |
| | 2:21009501:G:T | missense | p.Ala2456Asp | -0.04 | 8.4E-01 | -0.20 | 4.1E-01 | 0.00 | 9.9E-01 | 0.05 | 8.2E-01 | 0.0022 | 1.7 | Inf | 0.1 | 0.1 | 1.0 | Inf | 0.0 |
| | 2:21015135:G:T | missense | p.Leu1212Met | -0.04 | 8.6E-01 | -0.04 | 8.7E-01 | 0.07 | 8.1E-01 | -0.03 | 9.2E-01 | 0.0017 | 0.3 | Inf | 0.0 | 0.0 | 0.3 | Inf | 0.0 |
| *APOE* M4.1 | 19:44908847:C:A | missense | p.Ala184Asp | -0.86 | 1.3E-13 | -1.18 | 2.3E-20 | 0.02 | 8.7E-01 | -0.20 | 9.0E-02 | 0.0073 | Inf | NC | Inf | Inf | Inf | Inf | Inf |
| | 19:44908822:C:T | missense | p.Arg176Cys | -0.46 | 1.5E-03 | -0.69 | 2.4E-05 | -0.05 | 7.8E-01 | -0.04 | 7.8E-01 | 0.0049 | 0.0 | 0.1 | 0.0 | 0.1 | 0.0 | 0.0 | 0.0 |
| | 19:44908969:G:A | missense | p.Ala225Thr | 0.08 | 7.7E-01 | 0.13 | 6.6E-01 | -0.15 | 6.4E-01 | 0.08 | 7.8E-01 | 0.0011 | Variant not present in full American Indians | | | | | | |
| *PCSK9* M2.5 | 1:55052723:G:A | missense | p.Gly244Asp | -0.25 | 2.0E-04 | -0.46 | 4.7E-10 | 0.09 | 2.4E-01 | 0.18 | 1.0E-02 | 0.0237 | Inf | Inf | 2408.4 | Inf | Inf | Inf | Inf |
| | 1:55043951:G:A | missense | p.Gly106Arg | -0.59 | 6.3E-03 | -0.91 | 1.6E-04 | 0.12 | 6.4E-01 | -0.13 | 5.6E-01 | 0.0022 | Inf | Inf | Inf | Inf | Inf | Inf | Inf |

| | | | | | | | | | | | | | | | | | | | |
|---|---|---|---|---|---|---|---|---|---|---|---|---|---|---|---|---|---|---|---|
| | 1:55039995:C:T | missense | p.Ala53Val | -0.22 | 5.0E-02 | -0.25 | 4.1E-02 | 0.12 | 3.5E-01 | -0.14 | 2.1E-01 | 0.0083 | 0.0 | 0.0 | 0.0 | 0.1 | 0.0 | 0.0 | 0.1 |
| | 1:55043922:G:A | missense | p.Arg96His | -0.16 | 8.6E-02 | -0.22 | 4.4E-02 | 0.05 | 6.7E-01 | -0.11 | 2.6E-01 | 0.0114 | Inf | Inf | 1602.8 | Inf | Inf | 234.7 | Inf |
| | 1:55039974:G:T | missense | p.Arg46Leu | -0.45 | 1.3E-01 | -0.34 | 3.0E-01 | -0.67 | 5.7E-02 | 0.20 | 5.2E-01 | 0.0011 | 0.0 | Inf | 0.0 | 0.1 | 0.0 | Inf | 0.2 |
| | 1:55058129:A:G | missense | p.Asn425Ser | -0.41 | 1.6E-01 | -0.45 | 1.6E-01 | 0.08 | 8.0E-01 | 0.06 | 8.3E-01 | 0.0012 | Variant not present in full American Indians | | | | | | |
| | 1:55052746:G:A | missense | p.Val252Met | 0.38 | 1.7E-01 | 0.66 | 4.2E-02 | -0.42 | 2.0E-01 | 0.17 | 5.8E-01 | 0.0014 | 0.8 | Inf | 68.2 | Inf | 76.4 | Inf | 11.0 |
| | 1:55058549:C:T | missense | p.Arg469Trp | 0.28 | 3.0E-01 | 0.40 | 1.9E-01 | -0.66 | 4.2E-02 | 0.48 | 9.2E-02 | 0.0013 | 24.2 | Inf | 1.4 | 3.5 | 30.9 | Inf | 0.1 |
| | 1:55061388:C:G | missense | p.His565Gln | 0.21 | 3.5E-01 | 0.44 | 8.8E-02 | 0.00 | 9.9E-01 | -0.17 | 4.9E-01 | 0.0020 | 1.1 | Inf | Variant failed QC | | | | |
| | 1:55058550:G:A | missense | p.Arg469Gln | 0.11 | 6.8E-01 | 0.37 | 2.0E-01 | -0.04 | 9.0E-01 | -0.11 | 6.8E-01 | 0.0014 | Inf | Inf | 436.3 | Inf | Inf | 31.9 | Inf |
| | 1:55058182:G:A | missense | p.Ala443Thr | -0.05 | 7.9E-01 | -0.15 | 4.9E-01 | 0.16 | 4.9E-01 | 0.04 | 8.5E-01 | 0.0026 | 0.3 | 0.7 | 0.0 | 0.0 | 0.4 | 0.1 | 0.0 |
| *TM6SF2* M4.5 | 19:19269759:G:A | missense | p.Arg138Trp | -0.27 | 3.3E-08 | -0.20 | 1.2E-04 | 0.10 | 7.0E-02 | -0.33 | 2.5E-11 | 0.0461 | 2843.9 | Inf | 31.6 | 4.4 | Inf | 432.3 | Inf |
| | 19:19268740:C:T | missense | p.Glu167Lys | -0.23 | 1.4E-03 | -0.27 | 6.4E-04 | -0.06 | 4.7E-01 | -0.15 | 4.3E-02 | 0.0201 | 0.2 | 0.2 | 0.3 | 0.4 | 0.2 | 0.2 | 0.5 |
| | 19:19269704:A:G | missense | p.Leu156Pro | -0.08 | 7.5E-01 | 0.12 | 6.4E-01 | -0.09 | 7.6E-01 | -0.21 | 4.1E-01 | 0.0017 | 0.2 | 10.7 | 0.1 | 0.3 | 0.1 | Inf | 0.9 |
| *GPAM* M3.5 | 10:112159980:G:T | missense | p.Ser611Arg | 0.19 | 3.2E-03 | 0.11 | 1.4E-01 | 0.57 | 3.8E-14 | -0.17 | 9.0E-03 | 0.0250 | Inf | 383.1 | Inf | Inf | Inf | Inf | Inf |
| *LIPC* M4.1 | 15:58548359:G:A | missense | p.Glu280Lys | 0.01 | 9.5E-01 | -0.19 | 1.6E-01 | 0.38 | 9.4E-03 | 0.14 | 2.9E-01 | 0.0068 | 386.5 | Inf | 410.0 | 226.0 | 370.3 | 120.1 | Inf |
| | 15:58541878:A:G | missense | p.Ile123Val | -0.35 | 7.6E-02 | -0.35 | 1.1E-01 | 0.60 | 1.1E-02 | -0.62 | 2.3E-03 | 0.0026 | Inf | NC | Inf | Inf | Inf | Inf | Inf |
| | 15:58548541:C:G | missense | p.Phe340Leu | -0.07 | 7.6E-01 | -0.05 | 8.3E-01 | 0.24 | 3.5E-01 | -0.25 | 2.6E-01 | 0.0021 | 0.8 | Inf | Inf | Inf | Inf | Inf | Inf |
| *LIPG* M3.1 | 18:49582369:T:G | missense | p.His348Gln | 0.81 | 3.7E-03 | 0.62 | 4.8E-02 | 1.42 | 1.3E-05 | -0.21 | 4.7E-01 | 0.0013 | Inf | Inf | Inf | Inf | Inf | Inf | Inf |

[a] Variant is designated as chromosome:position:reference allele:alternate allele

[b] The unit is standard deviation of normalized traits.

[c] AAF: alternate allele frequency

The individual variants included in the masks of *APOB*, *APOE*, *PCSK9*, *TM6SF2*, *GPAM*, *LIPC*, and *LIPG* with ≥10 alternate allele counts were tested for association with plasma lipid levels in SWAI. The allele frequency ratios were derived by dividing the allele frequency in the full American Indians from the SWAI study by the allele frequencies in the European and East Asian individuals from DiscovEHR and TAICHI studies, and ancestral populations from the gnomAD exome database.

When variants were not captured and fall outside of the consistently covered region in a population, they were denoted as "not captured (NC)".
